# MultiCellDS: a standard and a community for sharing multicellular data

**DOI:** 10.1101/090696

**Authors:** Samuel H. Friedman, Alexander R. A. Anderson, David M. Bortz, Alexander G. Fletcher, Hermann B. Frieboes, Ahmadreza Ghaffarizadeh, David Robert Grimes, Andrea Hawkins-Daarud, Stefan Hoehme, Edwin F. Juarez, Carl Kesselman, Roeland M.H. Merks, Shannon M. Mumenthaler, Paul K. Newton, Kerri-Ann Norton, Rishi Rawat, Russell C. Rockne, Daniel Ruderman, Jacob Scott, Suzanne S. Sindi, Jessica L. Sparks, Kristin Swanson, David B. Agus, Paul Macklin

## Abstract

Cell biology is increasingly focused on cellular heterogeneity and multicellular systems. To make the fullest use of experimental, clinical, and computational efforts, we need standardized data formats, community-curated “public data libraries”, and tools to combine and analyze shared data. To address these needs, our multidisciplinary community created MultiCellDS (MultiCellular Data Standard): an extensible standard, a library of digital cell lines and tissue snapshots, and support software. With the help of experimentalists, clinicians, modelers, and data and library scientists, we can grow this seed into a community-owned ecosystem of shared data and tools, to the benefit of basic science, engineering, and human health.

## Unmet needs for collecting and curating multicellular data

Biology is increasingly focused on studying cellular heterogeneity and multicellular systems. Novel experiments, clinical trials, and simulation studies are generating incredible amounts of data on cell behavior, cell-cell and cell-matrix interactions, and cellular microenvironmental conditions. These advances are creating exciting new opportunities to formulate and test hypotheses, while synthesizing these disparate data sources to gain a deeper tissue-level understanding of health and disease.

However, the deluge of data has pushed existing data sharing and analysis paradigms to their limits. Key insights are effectively hidden in plain sight: tucked away in images, graphs, and tables; divorced from context; and inaccessible to computer analysis without significant manual work. While some data are online, much more are trapped offline on researchers’ flash drives, manually traded in emails, or inaccessible in private cloud storage. This severely limits data sharing, collaboration, and post-publication analyses that can offer new and unexpected insights.

There have been significant efforts to address these issues, but so far they have focused on describing genomic and molecular data (e.g., the Gene Ontology [1] for genetic data) or mathematical models (e.g., the Systems Biology Markup Language [2] for cell signaling models). None of these efforts have created a fixed data format for interchanging multicellular data or collected cell phenotype insights from many labs into shared, community-curated libraries with a uniform format. And while vast troves of experimental and clinical image data are available online to drive machine learning, we lack a standardized way to record extracted features, such as cell positions, sizes, shapes, and immunohistochemical stain statuses. Moreover, our lack of standardized data prevents us from directly linking between experimental and computational model systems, while also hindering our efforts to reconcile experimental and simulation results against clinical knowledge. With standardized data, would could directly couple experimental, computational, and clinical workflows, develop unified tools, and exchange insights. This would be bring key aspects of information science to multicellular biology, while facilitating reproducibility. See **Figure. 1**.

To address these unmet needs, we developed the MultiCellular Data Standard (MultiCellDS), a community-driven project [3] that facilitates the quantitative recording of the cellular microenvironment and phenotype in the form of digital cell lines and digital snapshots. Community-curated, centralized repositories of standardized data will pave the way to new data workflows and pipelines that will revolutionize the way we collaborate and learn in multicellular biology. By sharing standardized data—with formats that work for experimental, clinical, and computational models systems—we can work together to bridge the knowledge divide between molecular cell biology and the phenotypes of multicellular systems, tissues, organisms, and patients.

**Figure 1:**
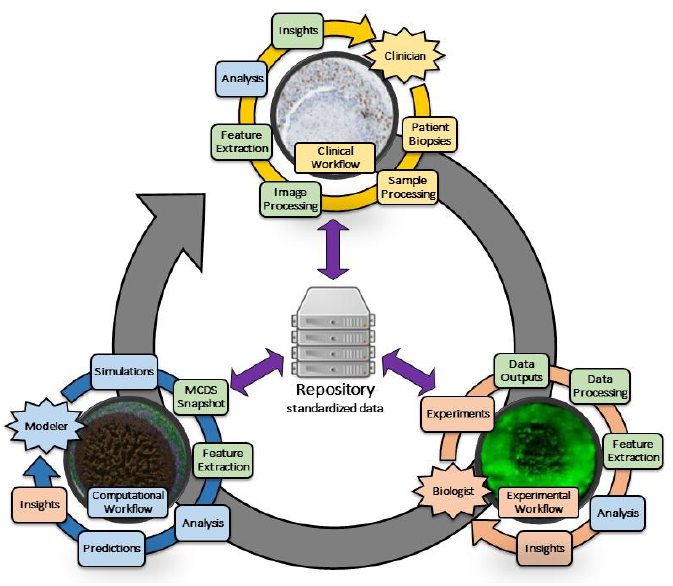
Integration of experimental, clinical, and computational workflows with standardized data. Standardized data formats make it possible to integrate experimental, clinical, and computational workflows, through shared data repositories and standards-compliant software tools. All three workflows share common tasks, such as image analysis, feature extraction, and data analysis. By adopting the same data formats, researchers in any workflow can leverage software tools and advances developed for the other workflows. Moreover, insights from each workflow could be shared in centralized repositories of shared data, potentially allowing new interactions between previously separate research teams and workflows.

### Box 1

#### Glossary of key terms and concepts in the MultiCellDS project

data element: a specific measurement (e.g., the radius of a cell nucleus); the smallest unit of data in MultiCellDS
XML: An extensible markup language (similar to HTML used for webpages), which can order data element hierarchically to match their relationships in biology
XML schema: A template for XML data, which describes the allowed data elements and how they can be arranged
metadata: Data about the data: extra information such as units, scale, uncertainty, or provenance
ontology: A controlled dictionary of allowed terms
OWL: A standardized ontology file, in the “web ontology language” format
provenance: The history of who created data, who revised it, who maintains it, and where it is published.
MultiCellDS: multicellular data standard
MultiCellDB: MultiCellDS database
MultiCellXML: MultiCellDS data, written in an XML format
phenotype dataset: A collection of cell phenotype and other measurements, in a single microenvironmental context
digital cell line: A collection of phenotype datasets and key metadata for a single biological cell line or type
digital snapshot: A readout of all cells, their phenotypes, and the microenvironment at a single time
collection: A logical grouping of digital cell lines, digital snapshots, collections, or a combination of these

## A community-driven project

To jump start the project, a core team—consisting of a cancer biologist, a mathematical modeler, data scientists, and a medical oncologist—drafted a working prototype of the data standard. To ensure that the standard could adapt to the diverse needs of the experimental, computational, and clinical communities, we assembled a group of invited “reviewers” (over thirty members spanning multiple disciplines, at institutions in the US and Europe) to critique the nascent standard and suggest improvements in three rounds of review. The invited reviews were supplemented by public talks and reviews to get feedback from the broader research community. The core team was responsible for leading the reviews, incorporating all feedback, and coordinating data and software contributions. This structure—a core team accountable to a skilled, multidisciplinary panel of reviewers—helped to balance the needs for extensive community feedback and involvement with the needs for fast and iterative development.

### Use cases to drive development

Each round of review was driven by a set of use cases: to represent cell phenotype measurements as *digital cell lines* (round 1); to record simulation and experimental data as *digital snapshots* (round 2); and to record segmented pathology data and de-identified clinical annotations in digital snapshots (round 3). These terms are defined in Box 1 and the descriptions below. Each round of review iteratively refined the data standard until we could complete the use cases. This helped ensure that the standard was not just a theoretical dictionary of terms, but a workable data language. Tackling unexpected problems suggested new data elements and metadata that could never have been anticipated purely through brainstorming and committee meetings. As a side effect, this process helped populate an initial “public library” of data, while driving software development for data analysis, visualization, and simulation.

## Digital snapshots: flash-freezing multiscale biology

Biological systems are typically observed or simulated at discrete, sampled times. Digital snapshots allow us to “freeze” these systems at a single time point and systematically record their states. Each snapshot begins with metadata: information on who generated the data and how (provenance) and other relevant details. A snapshot then records the microenvironmental context (e.g., oxygen concentration), either spatially or as average values. The snapshot closes with the multicellular data: e.g. cell positions, their phenotypes (at scales ranging from receptor status to gross behavior and morphology), and their types, if known (Figure 2). We use the same data elements for *in vitro, in vivo*, and *in silico* systems to facilitate interdisciplinary work.

**Figure 2:**
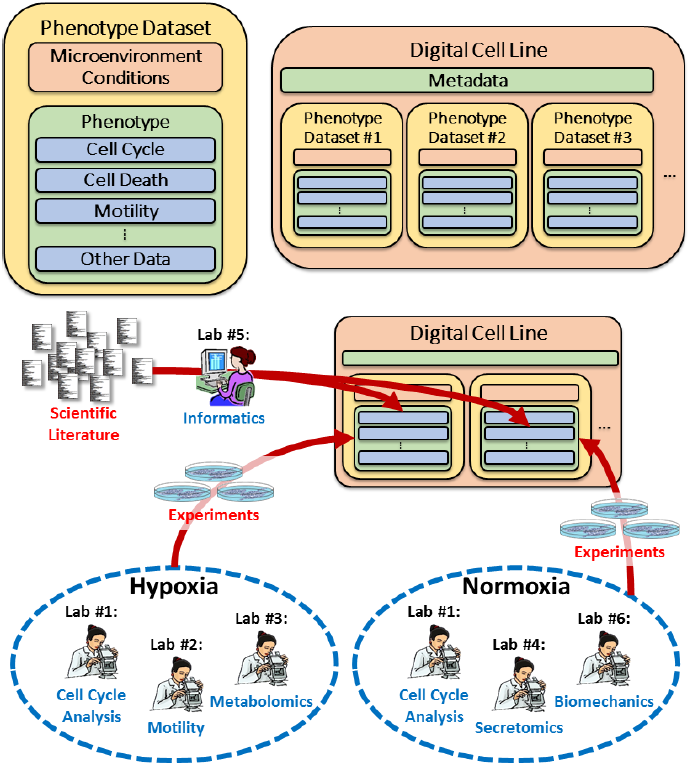
Digital cell line overview. <u>Top left:</u> Main elements in a *phenotype dataset* include microenvironmental conditions (context) and one or more phenotype measurements, grouped by function. <u>Top right:</u> A *digital cell line* collects metadata and multiple phenotype datasets. <u>Bottom:</u> Many labs can contribute to a digital cell line according to their expertise, while active community curation can mine the scientific literature to fill in the gaps in knowledge. Over time, digital cell lines aggregate biological insights from many sources.

## Digital cell lines: putting cell phenotype in context

By analyzing time series of digital snapshots, we can quantitate cell phenotype and correlate it with microenvironmental conditions. A digital cell line collects such measurements for a single cell type as phenotype datasets, allowing systematic recording of cell behavior in a single microenvironmental context (e.g., under normoxic culture conditions). To better systemize our current knowledge while exposing missing data, we cluster cell phenotype in several functional groups: cell cycling, cell death, mechanics, adhesion, motility, pharmacodynamics, secretion and uptake processes, and cell size/mass/morphology. A digital cell line can contain many phenotype datasets if it has been studied in many conditions, and each phenotype dataset can expand as our knowledge increases. Each phenotype dataset is matched to a description of the microenvironmental context, and it can be extended to embed any scale of data, such as “omics” data (Figure 3).

**Figure 3:**
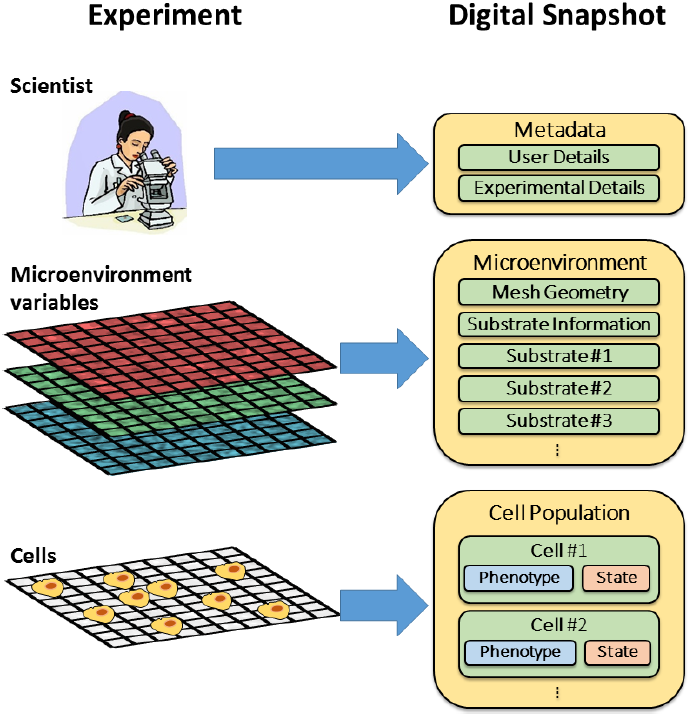
Digital snapshot overview. A digital snapshot captures <u>metadata</u> including details on the scientists (user metadata), cell lines and growth medium used (experimental details), information on the <u>microenvironment</u> such as one or more substrates sampled on a mesh, and a list of <u>cells</u> including their positions (state) and phenotypes. This storage paradigm can be applied to simulation data and segmented pathology images.

### Box 2

#### Versioning, Collections, and curation

To help with curation and reproducibility, digital cell lines are maintained with unique hierarchical classification numbers that increase as the data are changed. Drawing upon good practices from version control software in software engineering, digital cell lines can be split (forked) into branches maintained by different groups with different strengths and interests, and subsequently re-merged into a master branch to incorporate all the improvements. Related digital cell lines (or snapshots) can be bundled into collections to communicate ideas or knowledge that span multiple digital cell lines and/or snapshots, such as time series data (many digital snapshots) or patient cohorts (many patient-derived digital cell lines and corresponding digital pathology snapshots). These features help MultiCellDS to represent the variety and multitude of data expected in multicellular experimental, simulation, and clinical biology.

## A public library of digital cell lines and tissue snapshots

While testing and refining the data standard, we nucleated a “public library” of open data. To test whether digital cell lines could work beyond human cancer cell lines, we created digital cell lines for murine lymphoma, endothelial cells (to demonstrate highly motile, non-cancerous cells), yeast (our first non-mammalian cells), and bacteria (our first prokaryotic cell lines). Beyond “standardized” cell lines like MCF-7, we also created patient-derived digital cell lines for glioblastoma multiforme [4] and ductal carcinoma *in situ* of the breast [5]. In the course of creating over 200 digital cell lines, we demonstrated that the hierarchical phenotype dataset structure could scale from basic (e.g., parameters derived from ATCC culture protocol documents [6]) to extremely detailed (e.g., MCF-10A and MDA-MB-231 lines derived from a multi-institution study [7]). We also seeded the library with digital snapshots, including reference cancer simulation datasets from [8] and [9] and segmented breast cancer pathology images [10], including patient clinical annotations. Over the next several years, we plan to drastically extend this public library to include segmented TCGA pathology data [11] and segmented mouse liver data [12]. This entire library—stored in a centralized repository called MultiCellDB (multicellular database; see http://portals.MultiCellDS.org)—is freely available under the CC BY 4.0 license.

## Incentivizing good behavior: Rewarding contribution via attribution

There are three major types of contributions for community-curated data in MultiCellDB: generating the original data or measurements; performing data analysis or transformation; and actively curating the data (potentially from many sources!). All three types of contributions are essential, and they should be tracked in the provenance for reproducibility, transparency, and proper attribution. Moreover, the software tools used for data analyses and transformations need to be properly recorded. When a digital cell line or snapshot is used in a later publication, it is essential to record this chain of contributions, not only for reproducibility, but also to incentivize future contributions. Succinctly citing a chain of contributions is challenging, but we propose the following form:

> “We used digital cell line MCF-7 *[refs1]* version *n1* (MultiCellDB *id1*), created with data and contributions from [*refs2, refs3*].”

Here, *refs1* cites the publications or preprints that created the first and current version of the digital cell line, *refs2* cites the original data source(s), and *refs3* cites tools, software, and post-publication analyses or protocols used to transform the original data *(refs2)* into MultiCellDS data elements. It is also important to cite a fixed version of the digital cell to ensure that future replication studies use the same data. These are recorded as the version number (n1) and a unique identifier *(id1)*. (Box 2). Additional details on provenance and other metadata tracking can be found in the further resource documents below.

## The payoffs for sharing data in a common format

There are additional benefits to having a common format for multicellular data. With a fixed target of data elements, software developers across labs can work together to write data analysis, visualization, and simulation software that can be connected into sophisticated, reproducible research pipelines (Figure 4). This should lead to higher-quality software with development costs spread over more labs, while allowing researchers to cross-validate their results in a variety of compatible tools.

Because MultiCellDS uses the same data elements for experimental, clinical and simulation data, we can even use the same tools across disciplines, allowing better integration of experimental data into simulations, and more quantitative model validation. Insights in one domain can more readily “cross-pollinate” advances in the others when the data can be seamlessly read by the same tools across disciplines. Storing standardized data in centralized data repositories helps to archive critical data in the long term, thus improving reproducibility and repeatability. Moreover, MultiCellDS contributors can potentially widen their impact with increased data reuse and citations.

**Figure 4:**
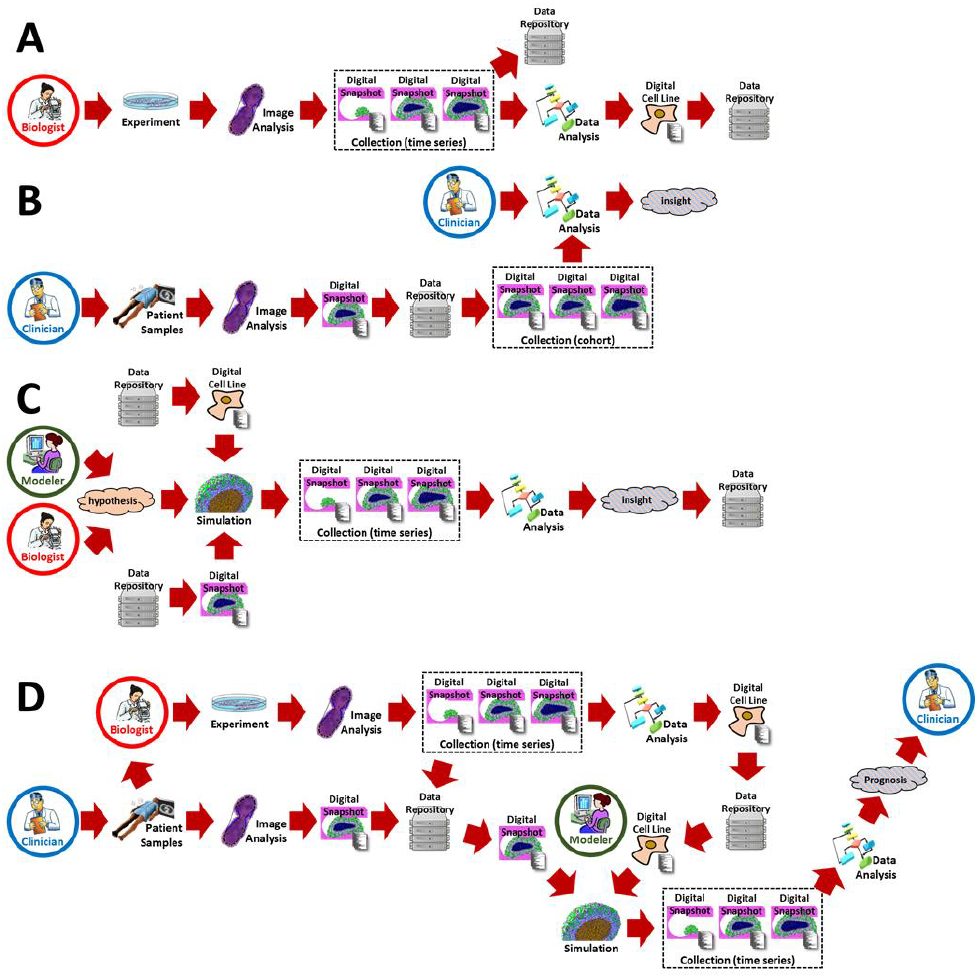
Sample integrated workflows using MultiCellDS data: Using standardized MultiCellDS data can help us to integrate experimental, clinical, and computational data in multidisciplinary workflows with unified software. **A:** A <u>biologist</u> uses standard image analysis software to save experimental images as digital snapshots, and then applies standard data analysis software to the snapshots to get new insights. New insights are shared in the data repository. **B:** A <u>clinician</u> uses standard image analysis software to save patient imaging as digital snapshots. Standard data analysis tools are applied to the digitized patient cohorts to get new insights. The insights are shared in the common repository. **C:** A <u>modeler</u> and a <u>biologist</u> form hypotheses to create a mathematical model, and they download digital snapshots and cell lines from the repository to seed the simulation. They analyze the simulation snapshots to gain new insights. **D:** A <u>clinician</u> analyzes a clinical sample as before. A <u>biologist</u> grows a patient-derived cell line from the clinical sample, performs experiments, analyzes the data, and creates a patient-specific digital cell line. A <u>modeler</u> uses the patient-de-rived data to simulate tumor progression under therapy, and sends the predicted prognosis back to the <u>clinician</u>.

### Getting a bird’s eye view of biology with centralized data repositories

By uniformly collecting cell phenotype knowledge in a centralized repository (MultiCellDB), we gain a unique opportunity to take a step back from focused single-lab investigations to compare cell behavior across many cell types. This uniform recording will help us to identify conserved behavior as well as contradictory data that may point to unknown biology or experimental error. Moreover, we can more readily identify gaps in our knowledge, to more systematically plan future experiments.

## Future directions, challenges, and a call to arms

We have developed a standard to systematically record cell phenotypes, microenvironmental conditions, and the state of multicellular systems. In doing so, we created an initial “public library” of digital cell lines and tissue snapshots. Building upon this, the community is actively creating an ecosystem of standards-compliant software to analyze, visualize, and simulate these shared data. In the near future, multidisciplinary teams will mine the shared data repository to formulate new biological hypotheses, encode these into computer models, and compare simulation outputs to experimental and clinical data to test and refine the hypotheses. Standardized data and shared software tools could accelerate this process, helping to close the gap between the benchtop, mathematical models, and the clinic.

Yet challenges remain. Recording the phenotypes of many cells in snapshots—or of single cell types in digital cell lines—falls short of multicellular biology. This is analogous to many actors delivering monologues on a shared stage. Drama and biology get interesting when the players interact. We must next characterize cell *networks*, including cell-cell interactions, cell mutations, and cell lineages. These enhancements can be woven into MultiCellDS using complementary ontologies, such as the Cell Behavior Ontology [13].

We must also expand the phenotype datasets to incorporate other critical measurements, such as genomic, proteomic, and metabolomics data. Moreover, we must account for the hysteresis in cell phenotype parameters: cells undergoing stresses do not always return to their original phenotypes once the stresses are removed. The community has begun testing ideas to addresses these key facets to multicellular systems biology, but broader participation is needed.

This early groundwork relied heavily upon data scientists and mathematicians to define data elements to characterize cell phenotype, microenvironmental conditions, and metadata. It is time to grow the community. We need experts across experimental biology and clinical practice to improve and expand the digital cell lines. We need a community discussion on what it means to improve a measurement—how do we know that a new measurement of cell motility is better than the old one? This will lead to useful discussions of not just reproducibility, but quality assurance in experimental pipelines. Over time, it will provide a useful library of “reference phenotype values” to help experimentalists to quantitatively compare their findings and separate protocol differences from genuine biological effects.

Moving forward, we must ensure that data donation and curation are user-friendly, and that the data standard evolves to meet the needs of the community. Just as community encyclopedias rely upon volunteer editors to update articles, we need volunteers to transform the scattered treasure of unannotated data into curated, standardized data in MultiCellDB. As MultiCellDS grows, we anticipate making the leap from a grassroots effort to a self-sustaining community, where the availability of standardized data and compatible software drives further adoption, contributions, new techniques, and community growth. Lastly, it is up to the community to make use of the data: to contribute more data; to mine the data for patterns that drive new hypotheses; to test new hypotheses in computational, experimental, and clinical models; and to unlock new knowledge that drives scientific progress and yields new therapies and strategies to improve health.

## Acknowledgements

We thank the Breast Cancer Research Foundation for the generous support in developing the MultiCellDS standard and initial repository. This work was also funded in part by the National Cancer Institute (5U54CA143907, 1R01CA180149). We thank the Lawrence J. Ellison Institute for Transformative Medicine for institutional support. Thanks also to the MultiCellDS review panel and the broader scientific community for their encouragement, participation, and data and software donations. Without a community, a data standard is just an XML file.

## Further resources

The MultiCellDS project website is hosted at http://MultiCellDS.org. A list of MultiCellDB portals, including the MultiCellDB reference repository, can be found at http://portals.MultiCellDS.org.

Extra documentation on MultiCellDS includes:

1. A more detailed “white paper” on the MultiCellDS standard and development process. [http://dx.doi.org/10.1101/090456]
2. User-focused overview of the data standard. [https://dx.doi.org/10.6084/m9.figshare.4269254.v1].
3. List of supported cell cycle representations. [https://dx.doi.org/10.6084/m9.figshare.4269263.v1]
4. Computer-generated documentation on the full standard, based upon the XML schema. [https://dx.doi.org/10.6084/m9.figshare.4269269].
5. The XML schema that official encodes the data standard. [https://dx.doi.org/10.6084/m9.figshare.4269272.v1 and https://dx.doi.org/10.6084/m9.figshare.4269275].
6. OWL ontology. [http://MultiCellDS.org/ont/multicellds.owl].
7. A protocol to transform DCIS pathology data into patient-derived digital cell lines. [https://dx.doi.org/10.6084/m9.figshare.4269248.v1].
8. Mathematical models used in the Chaste demonstration of MultiCellDS digital snapshots [https://dx.doi.org/10.6084/m9.figshare.4272242].
9. Matlab script used to help create ATCC-based digital cell lines [https://sourceforge.net/proiects/multicellds/files/Tools/ATCC_to_digital_cell_lines/]
10. Community norms for curation, versioning, and new data elements [https://dx.doi.org/10.6084/m9.figshare.4272374.v1].
11. Current MultiCellDS invited reviewers. [http://MultiCellDS.org/Team.php#review_panel]
12. MultiCellDS invited reviewers (rounds 1-3, through Nov. 2016). [https://dx.doi.org/10.6084/m9.figshare.4272197]

All MultiCellDS documentation can be found at http://MultiCellDS.org/Documentation.php.

A list of MultiCellDS-compatible software is maintained at http://software.MultiCellDS.org.

## References

1. Ashburner, M., et al., Gene ontology: tool for the unification of biology. The Gene Ontology Consortium. Nat Genet, 2000. 25(1): p. 25–9.

2. Hucka, M., et al., The systems biology markup language (SBML): a medium for representation and exchange of biochemical network models. Bioinformatics, 2003. 19(4): p. 524–31.

3. Macklin, P. MultiCellDS Project Website. 2014-present; Available from: http://MultiCellDS.org.

4. Baldock, A.L., et al., From patient-specific mathematical neuro-oncology to precision medicine. Front Oncol, 2013. 3: p. 62.

5. Edgerton, M.E., et al., A novel, patient-specific mathematical pathology approach for assessment of surgical volume: application to ductal carcinoma in situ of the breast. Anal Cell Pathol (Amst), 2011. 34(5): p. 247–63.

6. Physical Sciences-Oncology Network. BACKGROUND INFORMATION AND SOPS. [cited 2015 May 25, 2015]; Available from: https://physics.cancer.gov/bioresources/SOPs.aspx.

7. Physical Sciences-Oncology Centers, N., et al., A physical sciences network characterization of non-tumorigenic and metastatic cells. Sci Rep, 2013. 3: p. 1449.

8. Mirams, G.R., et al., Chaste: an open source C++ library for computational physiology and biology. PLoS Comput Biol, 2013. 9(3): p. e1002970.

9. Macklin, P., et al., Patient-calibrated agent-based modelling of ductal carcinoma in situ (DCIS): from microscopic measurements to macroscopic predictions of clinical progression. J Theor Biol, 2012. 301: p. 122–40.

10. Dong, F., et al., Computational pathology to discriminate benign from malignant intraductal proliferations of the breast. PLoS One, 2014. 9(12): p. e114885.

11. Cancer Genome Atlas, N., Comprehensive molecular portraits of human breast tumours. Nature, 2012. 490(7418): p. 61–70.

12. Hoehme, S., et al., Prediction and validation of cell alignment along microvessels as order principle to restore tissue architecture in liver regeneration. Proc Natl Acad Sci U S A, 2010. 107(23): p. 10371–6.

13. Sluka, J.P., et al., The cell behavior ontology: describing the intrinsic biological behaviors of real and model cells seen as active agents. Bioinformatics, 2014. 30(16): p. 2367–74.

